# A three-dimensional Air-Liquid Interface Culture Model for the Study of Epstein-Barr virus Infection in the Nasopharynx

**DOI:** 10.1101/2020.08.31.272096

**Authors:** Phillip Ziegler, Yulong Bai, Yarong Tian, Sanna Abrahamsson, Anthony Green, John Moore, Stella E. Lee, Michael M. Myerburg, Hyun Jung Park, Ka-Wei Tang, Kathy H.Y. Shair

## Abstract

Epstein-Barr virus (EBV) infection is ubiquitous in humans and is associated with the cancer, nasopharyngeal carcinoma. EBV replicates in the differentiated layers of stratified keratinocytes but whether the other cell types of the airway epithelium are susceptible to EBV is unknown. Here, we demonstrate with primary nasopharyngeal cells grown at the air-liquid interface that the pseudostratified epithelium can be susceptible to EBV infection and we report that susceptible cell types with distinct EBV transcription profiles can be identified by single-cell RNA-sequencing. Although EBV infection in the nasopharynx has evaded detection in asymptomatic carriers, these findings demonstrate that EBV latent and lytic infection can occur in the cells of the nasopharyngeal epithelium.

## INTRODUCTION

Epstein-Barr virus (EBV) is a human tumor virus from the γ-herpesvirus family (*1*). Infection is chronic and mostly asymptomatic but in a subset of individuals, latent infection is associated with certain types of B-cell lymphomas and epithelial carcinomas (*1*). EBV-associated nasopharyngeal carcinoma (NPC) is endemic in Southeast Asians and Alaskan Inuits (*2*). Diet and host genetics are risk-factors for NPC but almost all NPC tumors share the characteristic of latent and clonal infection with EBV (*2*). Thus, EBV infection is not a passenger infection but precedes the expansion of the neoplastic cell. EBV immortalizes B-cells; however, there are no reports of immortalization in epithelial cells (*3, 4*). Accordingly, the molecular pathogenesis of EBV in epithelial cells has been enigmatic. In the absence of cancer, EBV-infected cells are rarely detected in the nasopharyngeal epithelium (*5, 6*). This infrequency may be due to robust immune surveillance and/or small areas of infection that are difficult to capture at any one time. Thus, studies on EBV molecular pathogenesis in the nasopharynx have relied on cell culture models. Conventionally, two-dimensional (2-D) culture is used to study EBV latent infection in epithelial cells but it does not recapitulate the many cell types of the three-dimensional (3-D) epithelium in the nasopharynx (*3*). Furthermore, EBV-infected cell lines in 2-D culture are largely refractory to reactivation (*3*). Both latent and lytic infection are thought to encourage the persistence and spread of EBV in the nasopharynx, which can predispose cells to neoplasia (*2*).

Differentiation-induced reactivation in oral stratified keratinocytes cultured in 3-D organotypic rafts explains the lytic pathology of EBV-associated oral hairy leukoplakia (*7, 8*). The molecular pathogenesis in the nasopharyngeal epithelium is less clear as experimental models of the human nasopharyngeal airway epithelium have not been developed for EBV (*3*). Other than stratified keratinocytes, almost half of the nasopharyngeal epithelium is composed of pseudostratified respiratory epithelium which consists of a variety of cell types (ciliated, mucosecretory, basal and suprabasal) (*9*). In this study, we present a *de novo* EBV infection model of the nasopharyngeal pseudostratified epithelium grown in 3-D. Nasal primary cell cultures are differentiated from conditionally reprogrammed cells of the human nasopharynx and grown in 3-D air-liquid interface (ALI) culture (*10-12*). To distinguish this type of pseudostratified ALI culture from other types of ALI culture with cell lines or organotypic rafts, we herein refer to the primary nasal cell pseudostratified ALI as “pseudo-ALI” culture. Conventionally, pseudo-ALI cultures of airway epithelial cells are used to study acute virus infections such as influenza virus (*13*), respiratory syncytial virus (*14*), other respiratory pathogens (*15*), and more recently SARS-CoV-2 (*16*). Here, we report that the pseudo-ALI culture can be applied to the study of a persistent γ-herpesviruses, EBV. Using primary cells from a collection of 9 donors, examples of both latent and lytic infection are observed. Evidence of variation in donor susceptibility is presented. We report on one of the first examples of EBV latent infection captured in primary nasopharyngeal cells. These latently-infected cells are only observed in select donors, suggesting that some individuals could harbour a latent, local reservoir.

## RESULTS

### Establishment of a 3-D pseudo-ALI model of *de novo* EBV infection

We have previously demonstrated that conventional ALI culture can reactivate EBV from the NPC cell line, HK1-EBV, to yield high infectious titres (>10^6^ infectious green Raji units per cm^2^) (*17*). To elucidate EBV pathogenesis in primary cells, a method was developed for *de novo* EBV infection in differentiated nasal epithelial cells in pseudo-ALI culture. Primary cells from the nasopharynx, at the site of the lymphoid-rich Fossa of Rosenmüller, were collected under direct visualization from adult immune-competent donors undergoing endoscopic nasal procedures for reasons other than cancer. Primary cells were expanded on irradiated mouse 3T3-J2 fibroblasts in the presence of ROCK inhibitor (Y-27632) and lifted to the air-liquid interface on collagen-coated transwell membranes for 4 weeks (*10, 18*). Once primary cells have differentiated into pseudo-ALI cultures, EBV inoculum was applied to the apical surface by co-culture with the EBV-positive Akata cell line reactivated with anti-human IgG. The producer Akata cell line is recombinantly-infected with EBV expressing neomycin resistance and the EGFP marker gene inserted into the non-essential *BXLF1*, herein referred to as rAkata (*19*). As mock control, target cells were co-cultured with EBV-negative Akata cells similarly treated with anti-human IgG antibody.

Cells differentiated in pseudo-ALI culture were analyzed by histopathology to control for differentiation into respiratory epithelium (Supplementary Fig. 1). Hematoxylin and eosin stain demonstrated the presence of pseudostratified epithelium and ciliated cells. Alcian blue and periodic acid Schiff stain revealed mucin-secreting cells.

Immunohistochemistry staining for the proliferation marker, Ki67, showed the infrequent presence of cycling cells in the basal layer. To identify susceptible samples, EBV molecular diagnostics for latent and lytic markers of infection were developed for whole-mount staining of pseudo-ALI culture. These molecular diagnostics were first validated in the HK1-EBV cell line, in which 2-D culture is strictly latent but 3-D ALI culture triggers lytic reactivation. The detection of EBV-encoded RNAs (EBERs) by *in situ* hybridization (EBER-ISH) in the nucleus identifies latently-infected cells, while immunofluorescence staining for Zebra (immediate-early protein) in the nucleus or gp350 (late glycoprotein) in the cytoplasm denotes lytic infection (Supplementary Fig. 2). Notably, EBERs are not detected by EBER-ISH in oral hairy leukoplakia, a permissive EBV infection, and are thus a diagnostic marker of latent infection (*20*). Staining for the EBV oncoprotein, LMP1, identifies both latent and lytic infection. Stained images are scored by pixel intensity represented as a histogram compared to the mock (Fig. 1). Punctate LMP1 foci can also be discriminated as particles and scored for particle intensity, represented as a box and whisker plot (Fig. 1C).

**Figure 1.**
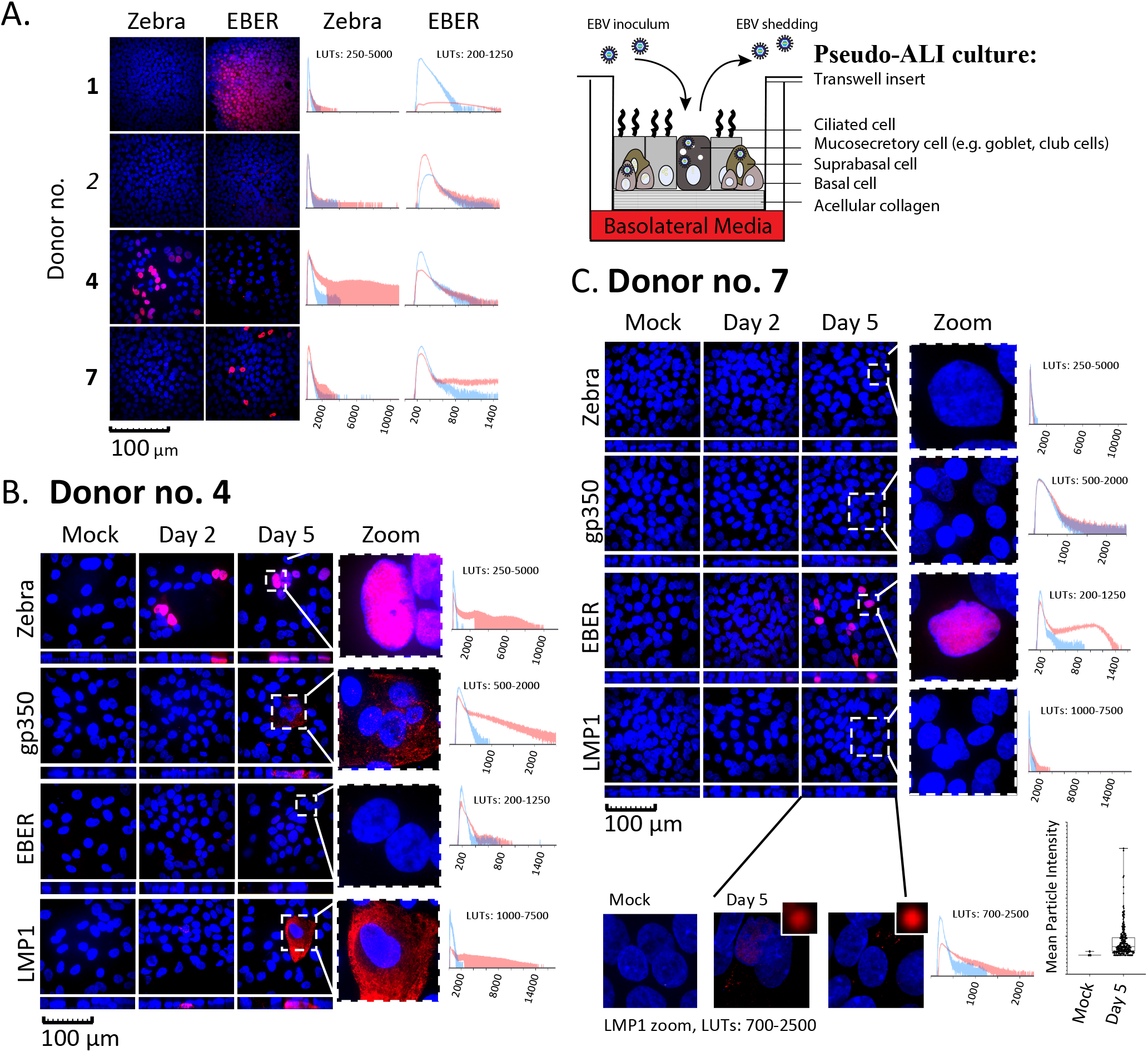
EBV *de novo* infection of primary nasopharynx-derived epithelial cells in pseudo-ALI culture. **A-C**, Immunofluorescence staining and EBER-ISH images (red) for EBV molecular diagnostics in pseudo-ALI cultures infected with EBV-positive rAkata B-cells or EBV-negative Akata B-cells (mock), counterstained with DAPI (blue). Shown are maximum intensity projections of confocal images on the *xy* (square) and *xz* (rectangle) planes. The pixel signal intensity of the EBV markers in the unzoomed image, in mock (blue line) and EBV-infected (red line) samples, are compared in the histograms. **(A)** Shown are the results for four donors. Cells were harvested at days 4-5 p.i. for Zebra and days 5-7 p.i. for EBER-ISH. More extensive analysis at days 2 and 5 p.i. was performed for donors **(B)** no. 4 and **(C)** no. 7. LMP1 foci in the weakly stained sample (donor no. 7) was also scored as mean particle intensity, from the unzoomed image. Bold, susceptible donor sample; italics, non-susceptible donor sample. LUTs, look-up-tables.

### EBV infection in pseudo-ALI culture show variation in donor susceptibility

Both susceptible and non-susceptible cultures were identified by EBV molecular diagnostics (Table 1). A total of 3 pseudo-ALI cultures (donor no. 1, 4, 7) were susceptible to EBV infection, while cultures from the other 6 donors were negative for the tested EBV molecular markers (Fig. 1A, Supplementary Fig. 3). Pseudo-ALI cultures from 2 donors (nos. 1 and 7) were positive for markers of latent infection, while cultures from donor no. 4 were positive for markers of lytic infection (Fig. 1A-C, Table 1). Stitched images showed no evidence of residual B-cell contamination from the inoculum after immunostain processing (Supplementary Fig. 4). In some cases, susceptible and non-susceptible cultures could be identified in the same experiment using the same stock of reactivated inoculum (Table 1). Thus, a failure to infect was indicative of host variation. Infections were repeated on low-passaged cells thawed from banked conditionally reprogrammed cells. In almost all cases of biological repeats (53 out of 54), either from susceptible (donor no. 4 and 7) or non-susceptible (donor no. 3, 5 and 8) donors, the same result in susceptibility and latent/lytic profiles were observed (Table 1, parentheses). Susceptibility to EBV did not appear to correlate with the presence or absence of comorbidity, although the number of samples collected is too small for statistical analysis. The EBV entry receptor for epithelial cells, ephrin receptor A2 (EphA2) (*21, 22*), was detected on the plasma membrane in all susceptible pseudo-ALI cultures but some of the cultures from non-susceptible samples were also strongly positive for EphA2 staining, such as donors no. 3 and 6 (Supplementary Fig. 1, Table 1). This indicates that while expression of EphA2 is consistent with EBV infection, other host factors also dictate susceptibility.

**Table 1.** Summary of molecular diagnostic results from EBV infection of nasal pseudo-ALIs with rAkata.

| Donor No.*<br>(reference code) | Experiment<br>No. | Susceptibility<br>to EBV<br>(-, +) | EphA2** | Latent | Lytic<br>(immediate-early, late) |  | Latent-<br>lytic | Reason for surgery/<br>comorbidity [other] |
| --- | --- | --- | --- | --- | --- | --- | --- | --- |
|  |  |  |  | EBER<br>(-/+) | Zebra<br>(-/+) | gp350<br>(-/+) | LMP1<br>(-/+) |  |
| 1 (H04) | 1 | + | ++ | + | - | n/a | n/a | CRSsNP***/none |
| 2 (H05) |  | - | + | - | - | n/a | n/a | Fungus ball, septal deviation,<br>allergic rhinitis/inflammatory<br>polyarthropathy |
| 3 (H08) | 2 | - | +++ | - (6/6) | - (6/6) | n/a | n/a | CRSsNP/inflammatory<br>polyarthropathy, Yellow Nail<br>Syndrome |
| 4 (H10) | 3 | + | ++ | - | + (5/6) | + | + | Left turbinate<br>hypertrophy/none |
| 5 (H12) | 4 | - | + | - (6/6) | - (6/6) | - | - | CRSsNP/inflammatory<br>polyarthropathy |
| 6 (H13) |  | - | +++ | - | - | - | - | Silent sinus syndrome/thyroid<br>disorder-unspecified, [Crohn's<br>disease] |
| 7 (H15) | 5 | + | ++ | + (6/6) | - (6/6) | - | +, weak | Septal perforation-unknown<br>etiology/[rheumatoid arthritis] |
| 8 (H17) | 6 | - | - | - (6/6) | - (6/6) | - | - | Septal deviation/none |
| 9 (H18) |  | - | + | - | - | - | - | Odontogenic sinusitis and<br>septal deviation/none |

\*Donor cells were obtained from the nasopharynx. The reference code denotes a unique donor identifier; \*\*Scoring criteria: -, +, ++, +++; \*\*\*CRSsNP, Chronic rhinosinusitis without nasal polyps; n/a, not available; biological repeats are indicated in parentheses.

### Molecular diagnosis of EBV infection reveals donor-specific differences in molecular pathogenesis – latent *vs*. lytic infection

Samples from donors no. 4 and 7 were subjected to more extensive analyses at days 2 and 5 post-infection (p.i.). Donor sample no. 4 stained positive for Zebra and LMP1 beginning at day 2 p.i., followed by gp350 at day 5 p.i., denoting a lytic infection (Fig. 1B). Donor sample no. 7 showed positivity for EBERs at day 5 p.i., denoting a latent infection (Fig. 1C). For donor sample no. 4, EBV replication was measured by qPCR of DNA harvested from extracellular or cell-associated DNase-resistant encapsidated virus (Supplementary Tables, A). As input control, pseudo-ALI cultures were fixed before co-culture with the inoculum. While the EBV genome copy number in the input control did not increase from day 2 to 5 p.i., extracellular EBV increased 37-fold (3.13 x 10^4^ copies at day 2 p.i. to 1.16 x 10^6^ copies at day 5 p.i.). EBV copy numbers did not increase in the cell-associated virus which measured between 1.55 – 4.06 x 10^4^ copies. This indicates that the majority of encapsidated EBV are packaged for secretion. Using virus collected from the extracellular source, infectious units were scored by the Green Raji Unit (GRU) assay in the non-producer Raji cell line. The secreted virus is indeed infectious, reaching 1.07 x 10^5^ GRUs by 5 days p.i. (Supplementary Tables, A).

### Single cell RNA-sequencing (scRNA-seq) reveals cell type-specific EBV transcriptional profiles

scRNA-seq analysis poses a challenge for all herpesvirus genomes because of overlapping 3’ co-terminal herpesvirus transcripts, whose non-uniquely mapped reads are discarded in the 10X Genomics single cell analysis pipeline (*23*). We reasoned however that this bioinformatics challenge is theoretically possible with the EBV gamma-herpesvirus genomes given that it has been demonstrated for alpha- and beta-herpesviruses (*24-26*). To identify EBV-infected cell types, donor sample no. 4 was subjected to scRNA-seq. Cell clusters (Fig. 2A), were assigned cell identities using experimentally-defined marker genes defined from primary human nasal epithelial cells grown in pseudo-ALI culture (*27*) and primary nasal tissue (The Human Cell Atlas Lung Consortium) (*28*), (Supplementary Fig. 5). All major airway epithelial cell types (basal, secretory, suprabasal and ciliated) could be identified (Fig. 2A). In order to improve alignment to the partially annotated EBV genome (NCBI KC207813.1), the reference genome for the Akata strain was updated with additional exon annotation totaling 87 genes. As there are no reports of scRNA-seq analysis on the EBV transcriptome, we tested several alignment algorithms using the 10X Genomics Cell Ranger pipeline. The reads were either aligned to the whole EBV genome as one annotation, as separately annotated genes, or as annotated genes but with genes that have regions of overlap in the same direction represented as fusion genes. Alignment to the separate annotation assigns the identity of EBV transcripts according to the reference annotations, but alignment to the other two annotations counts more EBV reads. Overall, the EBV transcriptome represents 0.08% (separate annotation) to 0.17% (one annotation and fused annotation) of the total transcriptome (Fig. 2B). This is similar to estimates from bulk RNA-seq of lymphoblastoid cell lines carrying latent EBV, where the majority of samples had EBV reads measuring 0.1-0.5% of the total transcriptome (*29*). A large majority of the cells (71%-82%) expressed EBV and/or EGFP transcripts (Fig. 2B).

**Figure 2.**
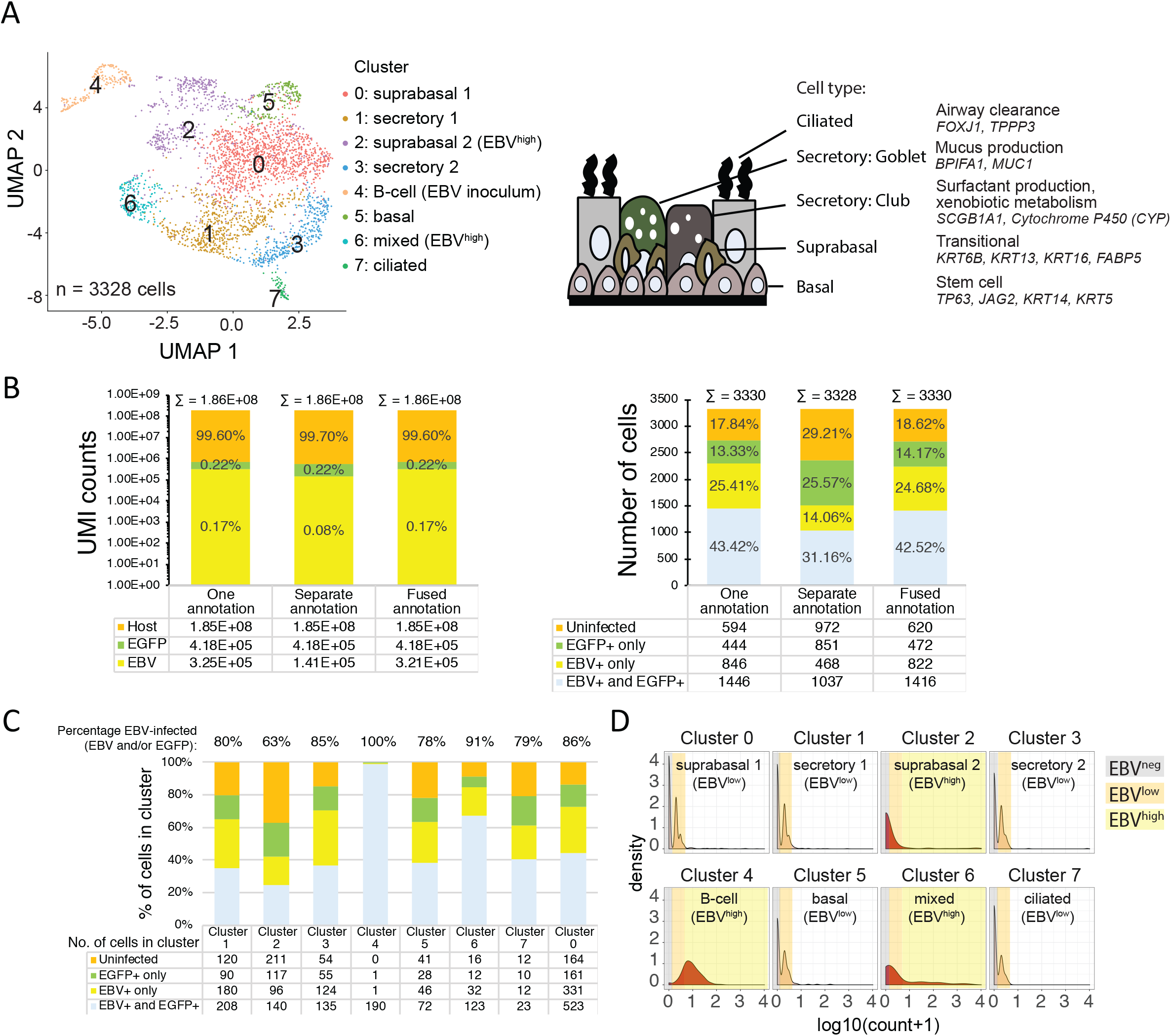
Identification of EBV-infected cell types from scRNA-seq performed on a pseudo-ALI culture from donor no. 4. (**A**) Uniform Manifold Approximation and Projection (UMAP) plots displaying the major cell clusters and assigned identity from host marker genes. Schematic displays the cell types in the pseudostratified nasal epithelium that are represented in the cell clusters. (**B**) Comparison of alignment methods against different EBV genome annotations, by UMI counts and numbers of cells. (**C**) The percentage of EBV-infected cells is displayed for each cluster. Shown are the results from alignment to the fused annotation EBV genome, (**D**) Density plots of log10 transformed pseudocount (UMI count+1) for EBV reads against normalized cell numbers displayed for each cell cluster. Cell numbers are normalized for the total number of cells in each cluster. Cells are grouped by EBV^negative^, EBV^low^ and EBV^high^ populations. Density plots with a similar profile are similarly coloured.

Every cluster scored positive for EBV and/or EGFP reads (Fig. 2C). *BHLF1, BHRF1, LF3 and LMP1/BNLF2a/BNLF2b* were the most frequently detected genes in the highest proportion of epithelial cells across clusters (Supplementary Fig. 6). All the cells in cluster 4 defined as the B-cell inoculum, expressed EBV and/or EGFP transcripts, with >97% of cells showing both EBV and EGFP (Fig. 2C). Across the epithelial cell clusters (clusters 1,2,3,5,6,7,0) the percent of cells with EBV reads ranged between 63%-91%, with no clear difference in susceptibility between clusters (Fig. 2C). However, density plots revealed two distinct EBV expression profiles, clusters with a peak at low UMI count (log_10_(count+1) < 0.3, clusters 0,1,3,5,7) denoted as EBV^low^, and clusters with 1-3 log_10_ higher UMI counts (clusters 2,4,6) denoted as EBV^high^ (Fig. 2D).

### Lytic infection is localized to suprabasal cells while latent infection is confined to basal/secretory and ciliated cell types

EBV^low^ cells found in all clusters displayed a distinct expression pattern (*BHLF1, BHRF1, LF3*, and the fused annotation *LMP-1/BNLFa/BNLFb*) which did not resemble a canonical type I/II/III latency profile (Supplementary Fig. 7). These cells are likely to be latent, refractory or in the early stage of the lytic cascade. These EBV^low^ cells are predominantly found in basal, secretory and ciliated cell types but also in a group of suprabasal cells defined by cluster 0 (Supplementary Fig.7). The mixed cell types in Cluster 6 could be further divided into 4 sub-populations with distinct marker gene expression (Supplementary Fig. 8A). EBV^high^ cells within sub-cluster 6-4 with basal cell features (Supplementary Fig. 8) and the EBV^high^ suprabasal cells in cluster 0 (Supplementary Fig. 7) display high levels of *BZLF1/BRLF1* indicative of reactivation. Lytic infection is mainly observed in EBV^high^ suprabasal cells (cluster 2) where there is global induction of EBV genes (Supplementary Fig. 7) but shut-off of host mRNA (Supplementary Fig. 9). Overall, all cell types in the pseudo-ALI were susceptible to EBV-infection; however productive virus infection is mainly confined to suprabasal cells. Despite the concerns of overlapping and low abundance viral transcripts evading capture by scRNA-seq (*23*), we conclude that with the appropriate reference annotation, the γ-herpesvirus genome can be analyzed by scRNA-seq.

## DISCUSSION

In conclusion, we demonstrate that pseudostratified epithelial cells from the nasopharynx are susceptible to EBV infection. Results from this study would indicate that host variables other than the expression of EphA2 impact susceptibility in the nasopharynx as well as the type of EBV infection (productive or non-productive). Given the relatively rare chance of finding an EBV-infected nasopharyngeal cell in asymptomatic carriers (*5, 6*), the pseudo-ALI culture thus provides a new organoid model in which to study EBV molecular pathogenesis in the nasopharynx. Our findings agree with prior studies in organotypic rafts using oral-derived keratinocytes that EBV lytic infection is confined to suprabasal cells (*8, 30*). We conclude that latent infection can occur in nasopharynx-derived basal/secretory/ciliated cell types, which may habour a local EBV reservoir, and suggest that the basal cell type could be a progenitor cell for NPC tumors.

## MATERIALS AND METHODS

### Samples

Primary nasopharyngeal cell samples were collected at UPMC Mercy hospital before emergence of the COVID-19 pandemic conducted under IRB STUDY#19030014 University of Pittsburgh Sinus Fluid and Tissue Bank. Voluntary informed consent was obtained for the collection, storage and analysis of biologic and/or genetic material for research, and such de-identified samples and de-identified data may be shared with other investigators for health research.

### Cell culture

The HK1 NPC cell line and the Akata Burkitt’s lymphoma B-cell line were maintained in RPMI supplemented with 10% fetal bovine serum. HK1 and Akata cells infected with the EBV recombinant Akata strain (courtesy of Dr. George Tsao, Hong Kong University) were supplemented with 800 μg/mL G418 selection(*19, 31*). The EBV-infected HK1 (HK1-EBV) and Akata (rAkata) cells express neomycin-resistance and EGFP from the SV40 early promoter, inserted into the EBV *BXLF1* locus, and are intact for expression of the EBV miRNAs(*19, 32*). Cells were incubated at 37°C with 5% CO_2_ and confirmed to be negative for mycoplasma contamination by PCR. Primary nasal epithelial cells were cultured from nasal cytobrush scrapings of the nasopharynx or inferior turbinate. Protocols for Collected cells were seeded on irradiated mouse 3T3-J2 feeder fibroblasts and expanded in Georgetown media(*10*). The presence of 4 µM ROCK inhibitor (Y-27632) extends the lifespan and induces the conditional reprogramming of epithelial cells(*18*). Media was changed daily, and cells were sub-cultured at 1:4 seeding density. At passage 1 or 2, 1.5×10^5^ cells were seeded on human type IV placental collagen-coated transwell filters (Corning, 0.33 cm^2^, 0.4 µm, polyethylene terephthalate) in Georgetown media for 24 hours. After 24 hours apical media was removed, cultures washed once in PBS, and the basolateral media was replaced with 400 µL of ALI medium(*11*) supplemented with 0.5% Ultroser G Serum Substitute (PALL), denoted as UNC/USG basolateral media. Cultures were maintained at the air-liquid interface for at least 4 weeks to allow differentiation into a pseudo-ALI culture. Basolateral media was changed 3 times a week. HK1 and HK1-EBV cells were cultured at the air-liquid interface as previously described(*17*).

### EBV Infection

rAkata EBV-infected cells was reactivated at 1×10^6^ cells/mL with a goat polyclonal anti-human IgG Fc-specific antibody (Sigma) for 48 hours. EBV-negative Akata cells were similarly treated with anti-human IgG antibody as a mock control. Virus production was confirmed by quantitative PCR for *BALF5*, as described in Supplementary Methods. Reactivated Akata cells were pelleted by centrifugation and resuspended at a concentration of 1.25×10^7^ cells /mL in calcium-/magnesium-free Dulbecco’s PBS (DPBS). Primary pseudo-ALI cultures were washed in DPBS once for 5 minutes at 37°C and twice briefly at room temperature. The reactivated B-cell suspension was added to the apical surface of the pseudo-ALI culture in 200 μL, basolateral media was replaced with DPBS, and cultures were pre-incubated at 37°C for 2 hours. The basolateral DPBS was then replaced with UNC/USG media and cultures incubated for a further 48 hours at 37°C. B-cell co-culture was removed by aspiration, and pseudo-ALI cultures were washed three times in Hank’s buffered saline solution (HBSS) to remove remaining B-cells. Cultures were fixed (2 days p.i.) or incubated at 37°C for up to 5 additional days (4-7 days p.i.), changing UNC/USG basolateral media every 48 hours.

### scRNA-seq

Cell suspensions were loaded into 10X Genomics Chromium instrument for library preparation as described previously (*33*), using the single cell 3’v3.1 (SC3Pv3) chemistry. Library QC was performed on an Agilent Bioanalyzer. High-throughput sequencing was performed by Novogene on a HiSeq paired-end 150 bp configuration yielding >472M reads.

### Code availability

The R script for Seurat workflow and for data visualization is available upon request.

### Data availability

Raw and processed scRNA-seq data files, and the merged EBV+hg38 genome annotation file will be deposited in NCBI Gene Expression Omnibus (GEO) GSE157243 upon publication. Filtering criteria and data processing steps are provided in the GEO submission. The Akata EBV genome (NCBI KC207813.1) with updated annotations are available in Github (https://github.com/TangLabGOT/Reference-Genomes).

This work is supported in part by the Hillman Foundation, the University of Pittsburgh Center for Research Computing, and used the Hillman Tissue and Research Pathology Services shared resource that is supported in part by National Institutes of Health award P30CA047904. We thank Tracy Tabib and Dr. Robert Lafyatis from the University of Pittsburgh Single Cell Core for advice on scRNA-seq and Dr. George S.W. Tsao (Hong Kong University, Hong Kong SAR) for providing the HK1 and HK1-EBV cell lines. We thank the Cystic Fibrosis Research Center cell core at the University of Pittsburgh (funded by the Cystic Fibrosis Foundation Research Development Program to the University of Pittsburgh) for providing reagents and advice on the culture of primary cells. We thank the Bioinformatics Core Facility at the Sahlgrenska Academy for bioinformatics support.

## AUTHOR CONTRIBUTIONS

K.S. designed experiments, analysed data and prepared the manuscript. P.Z. conducted experiments for EBV infection and molecular diagnosis. S.L. and J.M. collected specimens, comorbidity information and revised the manuscript. M.M. provided protocols and reagents for pseudo-ALI culture and revised the manuscript. P.Z., Y.B., Y.T., S.A., H.J.P. and K-W.T. analysed data and prepared the manuscript. A.G. established protocols and performed the histology on pseudo-ALI cultures.

## REFERENCES

1. L. S. Young, L. F. Yap, P. G. Murray, Epstein-Barr virus: more than 50 years old and still providing surprises. Nature reviews. Cancer 16, 789–802 (2016).

2. N. Raab-Traub, Nasopharyngeal Carcinoma: An Evolving Role for the Epstein-Barr Virus. Current topics in microbiology and immunology 390, 339–363 (2015).

3. K. H. Y. Shair, A. Reddy, V. S. Cooper, in Cancers (Basel). (2018), vol. 10.

4. S. W. Tsao et al., The biology of EBV infection in human epithelial cells. Seminars in cancer biology 22, 137–143 (2012).

5. R. Pathmanathan, U. Prasad, R. Sadler, K. Flynn, N. Raab-Traub, Clonal proliferations of cells infected with Epstein-Barr virus in preinvasive lesions related to nasopharyngeal carcinoma. The New England journal of medicine 333, 693–698 (1995).

6. C. K. Sam et al., Analysis of Epstein-Barr virus infection in nasopharyngeal biopsies from a group at high risk of nasopharyngeal carcinoma. International journal of cancer. Journal international du cancer 53, 957–962 (1993).

7. L. M. Hutt-Fletcher, The Long and Complicated Relationship between Epstein-Barr Virus and Epithelial Cells. Journal of virology 91, (2017).

8. R. M. Temple et al., Efficient replication of Epstein-Barr virus in stratified epithelium in vitro. Proceedings of the National Academy of Sciences of the United States of America, (2014).

9. M. Y. Ali, Histology of the human nasopharyngeal mucosa. J Anat 99, 657–672 (1965).

10. F. Serrano Castillo et al., A physiologically-motivated model of cystic fibrosis liquid and solute transport dynamics across primary human nasal epithelia. J Pharmacokinet Pharmacodyn 46, 457–472 (2019).

11. M. L. Fulcher, S. H. Randell, in Methods in molecular biology. (2013), vol. 945, pp. 109– 121.

12. S. Chen, J. Schoen, Air-liquid interface cell culture: From airway epithelium to the female reproductive tract. Reprod Domest Anim 54 Suppl 3, 38–45 (2019).

13. M. Richard et al., Influenza A viruses are transmitted via the air from the nasal respiratory epithelium of ferrets. Nature communications 11, 766 (2020).

14. C. S. Anderson et al., CX3CR1 as a respiratory syncytial virus receptor in pediatric human lung. Pediatr Res 87, 862–867 (2020).

15. W. Hao et al., Infection and propagation of human rhinovirus C in human airway epithelial cells. Journal of virology 86, 13524–13532 (2012).

16. R. Lu et al., Genomic characterisation and epidemiology of 2019 novel coronavirus: implications for virus origins and receptor binding. Lancet 395, 565–574 (2020).

17. E. A. Caves et al., Air-Liquid Interface Method To Study Epstein-Barr Virus Pathogenesis in Nasopharyngeal Epithelial Cells. mSphere 3, (2018).

18. X. Liu et al., ROCK inhibitor and feeder cells induce the conditional reprogramming of epithelial cells. The American journal of pathology 180, 599–607 (2012).

19. S. Maruo, L. Yang, K. Takada, Roles of Epstein-Barr virus glycoproteins gp350 and gp25 in the infection of human epithelial cells. The Journal of general virology 82, 2373–2383 (2001).

20. K. Gilligan, P. Rajadurai, L. Resnick, N. Raab-Traub, Epstein-Barr virus small nuclear RNAs are not expressed in permissively infected cells in AIDS-associated leukoplakia. Proceedings of the National Academy of Sciences of the United States of America 87, 8790–8794 (1990).

21. J. Chen et al., Ephrin receptor A2 is a functional entry receptor for Epstein-Barr virus. Nat Microbiol 3, 172–180 (2018).

22. H. Zhang et al., Ephrin receptor A2 is an epithelial cell receptor for Epstein-Barr virus entry. Nat Microbiol 3, 164–171 (2018).

23. D. P. Depledge, I. Mohr, A. C. Wilson, Going the Distance: Optimizing RNA-Seq Strategies for Transcriptomic Analysis of Complex Viral Genomes. Journal of virology 93, (2019).

24. N. Drayman, P. Patel, L. Vistain, S. Tay, HSV-1 single-cell analysis reveals the activation of anti-viral and developmental programs in distinct sub-populations. Elife 8, (2019).

25. E. Wyler et al., Single-cell RNA-sequencing of herpes simplex virus 1-infected cells connects NRF2 activation to an antiviral program. Nature communications 10, 4878 (2019).

26. M. Shnayder et al., Single cell analysis reveals human cytomegalovirus drives latently infected cells towards an anergic-like monocyte state. Elife 9, (2020).

27. S. Ruiz Garcia et al., Novel dynamics of human mucociliary differentiation revealed by single-cell RNA sequencing of nasal epithelial cultures. Development 146, (2019).

28. F. A. Vieira Braga et al., A cellular census of human lungs identifies novel cell states in health and in asthma. Nature medicine 25, 1153–1163 (2019).

29. A. Arvey et al., An atlas of the Epstein-Barr virus transcriptome and epigenome reveals host-virus regulatory interactions. Cell host & microbe 12, 233–245 (2012).

30. D. M. Nawandar et al., Differentiation-Dependent KLF4 Expression Promotes Lytic Epstein-Barr Virus Infection in Epithelial Cells. PLoS pathogens 11, e1005195 (2015).

31. A. K. Lo et al., Epstein-Barr virus infection alters cellular signal cascades in human nasopharyngeal epithelial cells. Neoplasia 8, 173–180 (2006).

32. A. K. Lo et al., Modulation of LMP1 protein expression by EBV-encoded microRNAs. Proceedings of the National Academy of Sciences of the United States of America 104, 16164–16169 (2007).

33. C. Morse et al., Proliferating SPP1/MERTK-expressing macrophages in idiopathic pulmonary fibrosis. Eur Respir J 54, (2019).

