## Supplementary Figures and Tables for "A three-dimensional Air-Liquid Interface Culture Model for the Study of Epstein-Barr virus Infection in the Nasopharynx"

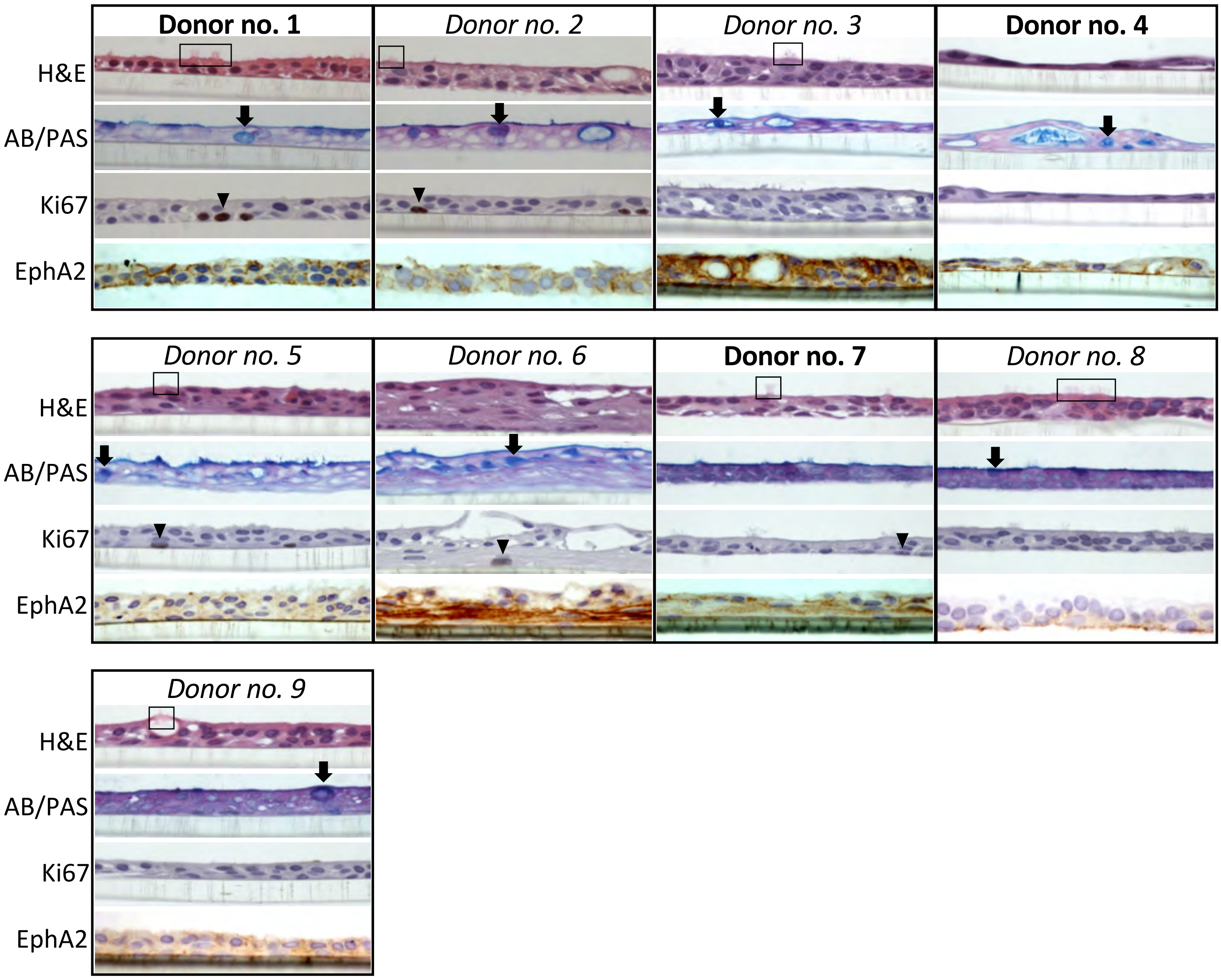


**Supplementary Figure 1: Histological analysis of pseudo-ALI cultures derived from the nasopharynx for markers of cellular differentiation and the EBV epithelial cell receptor.** Shown are pseudo-ALI cultures susceptible (bold) and non-susceptible (italics) to *de novo* EBV infection. Examples of ciliated cells are marked by a black box in the Hematoxylin & Eosin (H&E) stain. Mucin-producing cells are marked by a black arrow in the Alcian blue/periodic acid Schiff (AB/PAS) stain. Basal cells that are proliferating are marked by a black arrowhead in the Ki67 stain. Expression of the EBV epithelial cell receptor, Ephrin type-A receptor 2 (EphA2), is indicated by brown 3, 3'–diaminobenzidine (DAB) staining counterstained with nuclear blue. More extensive analysis at days 2 and 5 p.i. was performed for donors no. 4 and 7 (see Fig. 1).

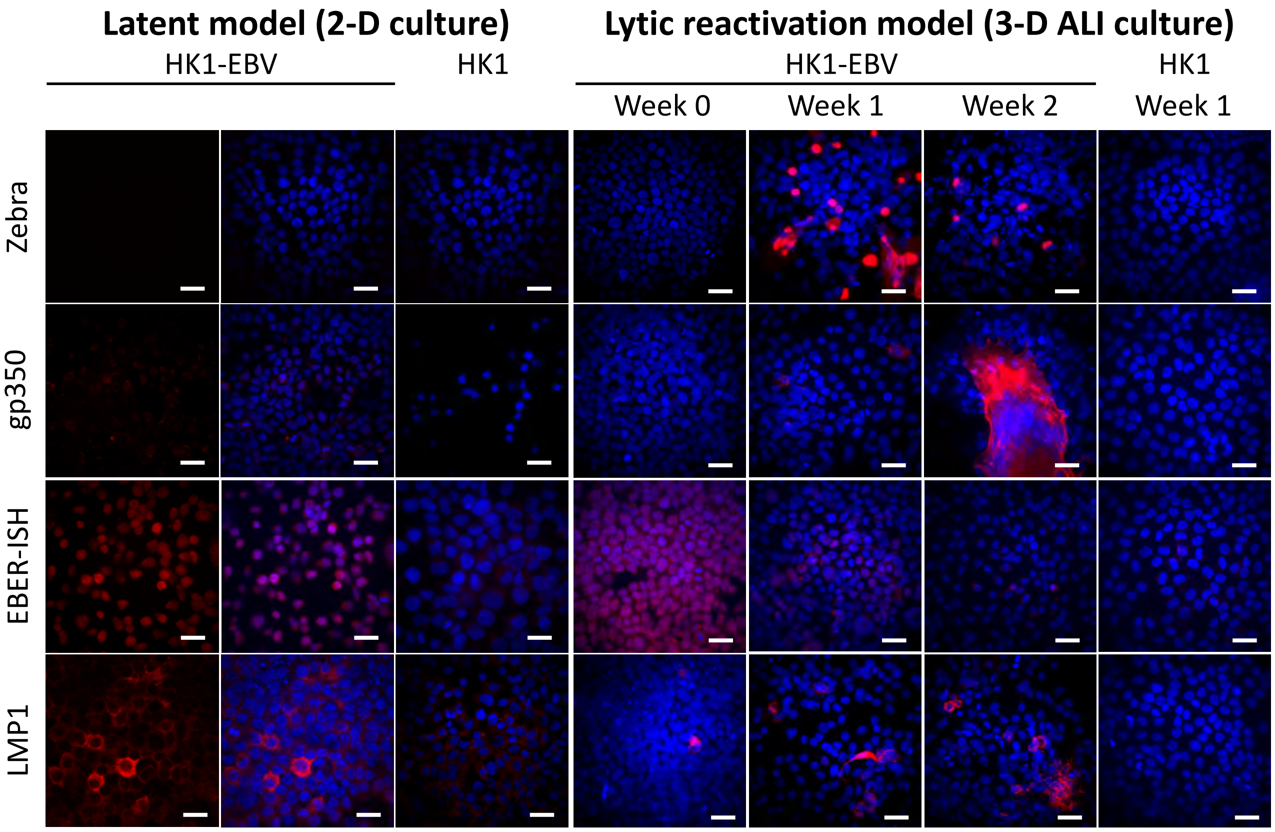


**Supplementary Figure 2. In-situ hybridization (EBER-ISH) and immunofluorescence staining (LMP1, Zebra, gp350) for EBV molecular markers in the HK1-EBV latent (2-D culture) and lytic reactivation (3-D ALI culture) cell culture model.** Shown are confocal images from one Z-section. Positive staining is indicated in red and nuclei are counterstained with DAPI (blue). Scale bar = 40 µm.

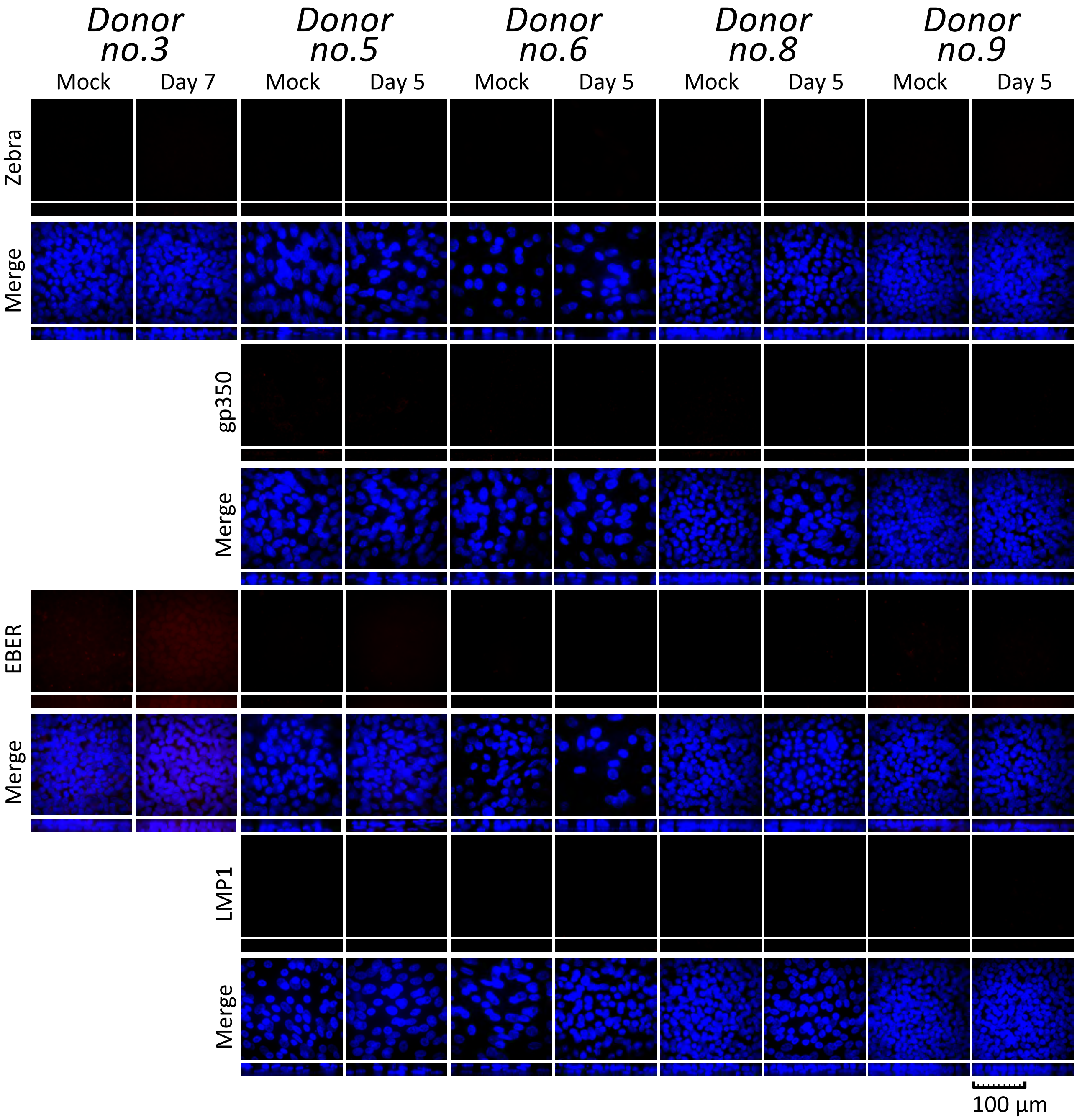


**Supplementary Figure 3. EBV *de novo* infection in non-susceptible pseudo-ALI cultures.** Shown are maximum intensity projections of confocal images on the *xy* (square) and *xz* (rectangle) planes. Nasopharyngeal cells in pseudo-ALI culture are stained for Zebra, gp350, LMP1, or EBER-ISH (red), and counterstained with DAPI (blue).

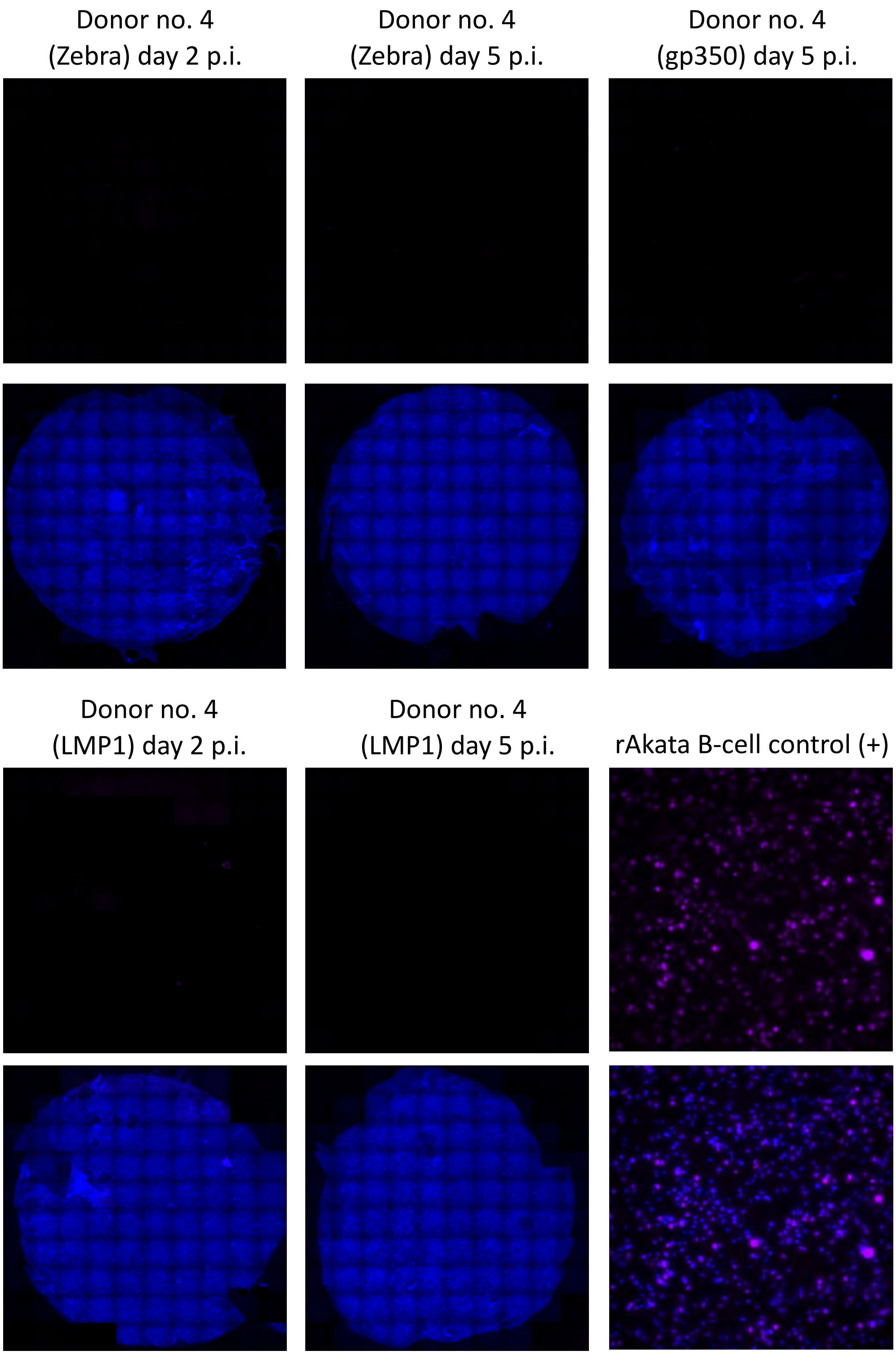


**Supplementary Figure 4. Anti-human IgG stains to control for contaminating B-cells from the inoculum.** Stitched images of pseudo-ALI cultures stained for anti-human IgG (purple) showing the entire membrane area, counterstained with DAPI (blue). Shown are examples of the control images for the corresponding stain (labeled in parentheses) in which positive staining for the EBV marker of interest was detected. The positive control (+) for anti-human IgG is an image of stained rAkata B-cells on a glass slide.

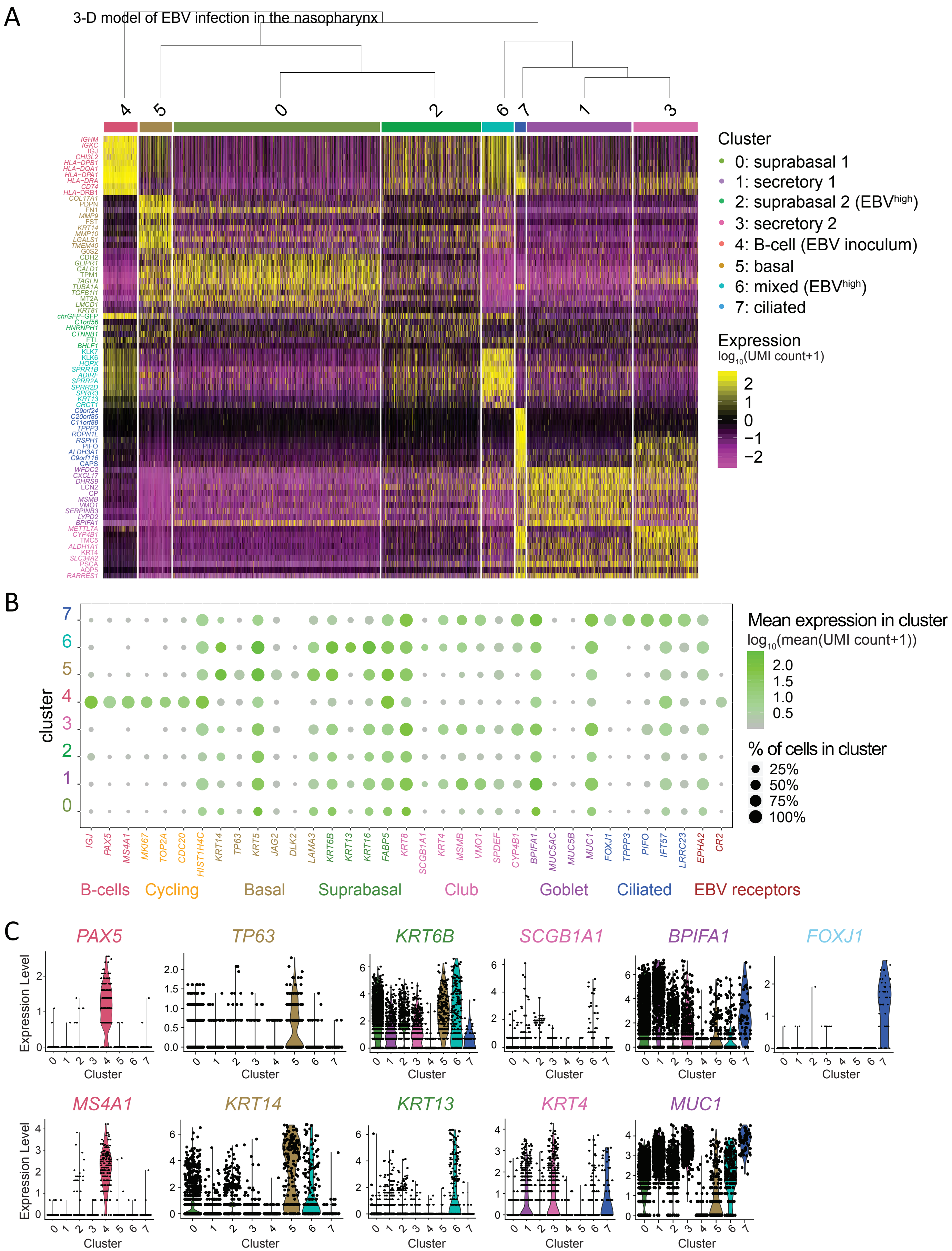


**Supplementary Figure 5. Assignment of cluster identity by cell type-specific marker genes for donor no. 4.** (A) Hierarchical clustering of heatmap representing cluster-specific top marker genes. Expression shows log10 transformed pseudocount (UMI count+1). (B) Dot plot representing marker gene expression by cluster. Dot size indicates the percentage of cells in each cluster expressing the marker gene. Colour gradient indicates the mean expression of each marker gene, averaged from positively scored cells, for each cluster. Cluster identity enriched for cell-type specific marker genes are colour coded. Cluster 6 expresses a mixture of marker genes indicative of multiple cell types (basal, suprabasal, goblet). The EBV receptor gene, *CR1*, is not shown as it was not expressed according to the count matrix. (C) A few of the highly cell type-specific marker genes were chosen for display by violin plots.

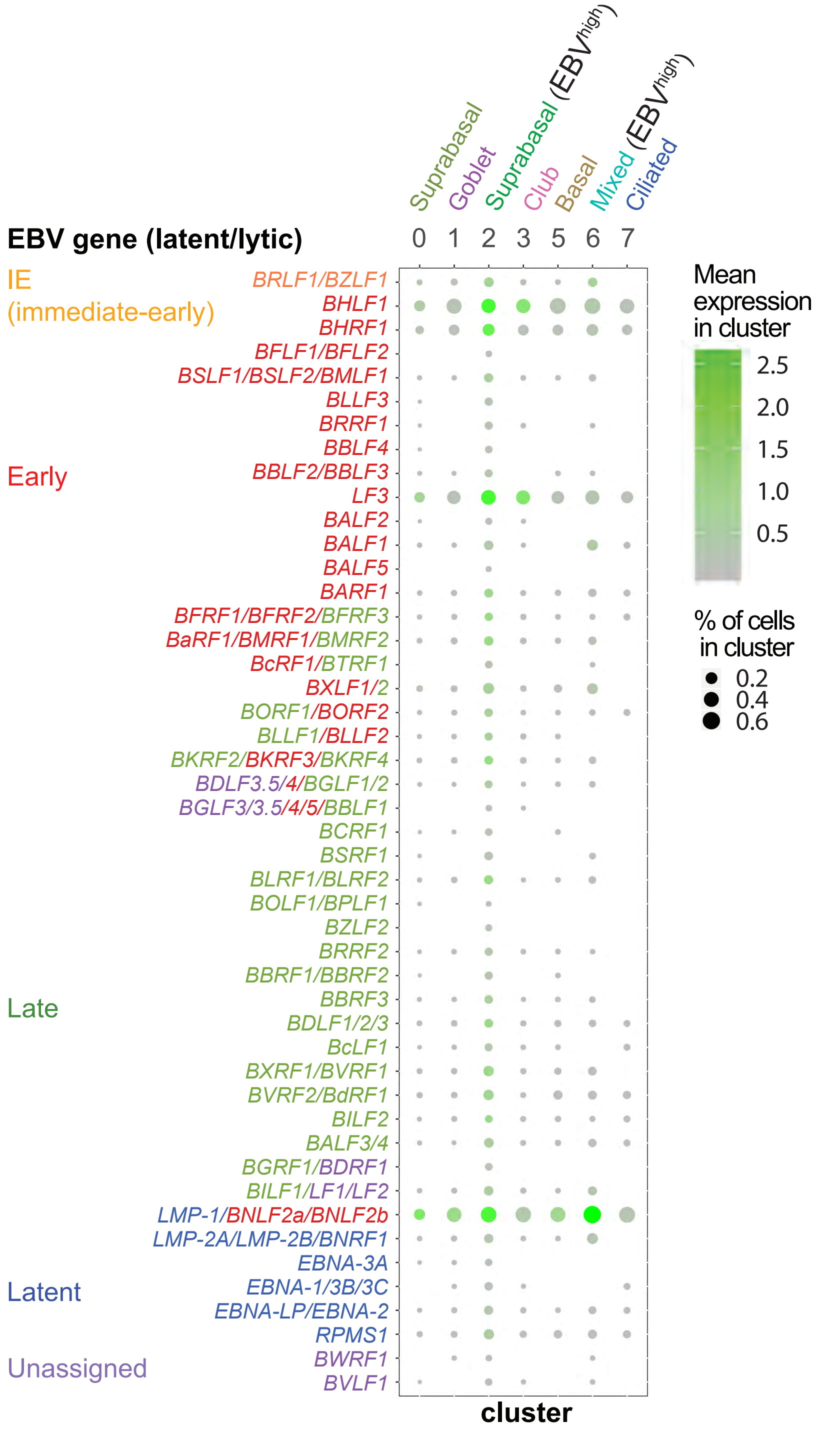


**Supplementary Figure 6. Averaged expression of EBV genes by cluster.** Dot plot shows expression of EBV genes for epithelial cell clusters in the EBV-infected pseudo-ALI culture of donor no. 4. Genes are grouped in latent or lytic (immediate-early/early/late). Unassigned genes are in purple. Mean expression by cluster shows log10 transformed pseudocount (UMI count+1) for EBV gene averaged by cluster. Dot size indicates the percentage of cells in each cluster expressing the EBV gene. The scRNA-seq reads were aligned to the EBV genome fused annotation.

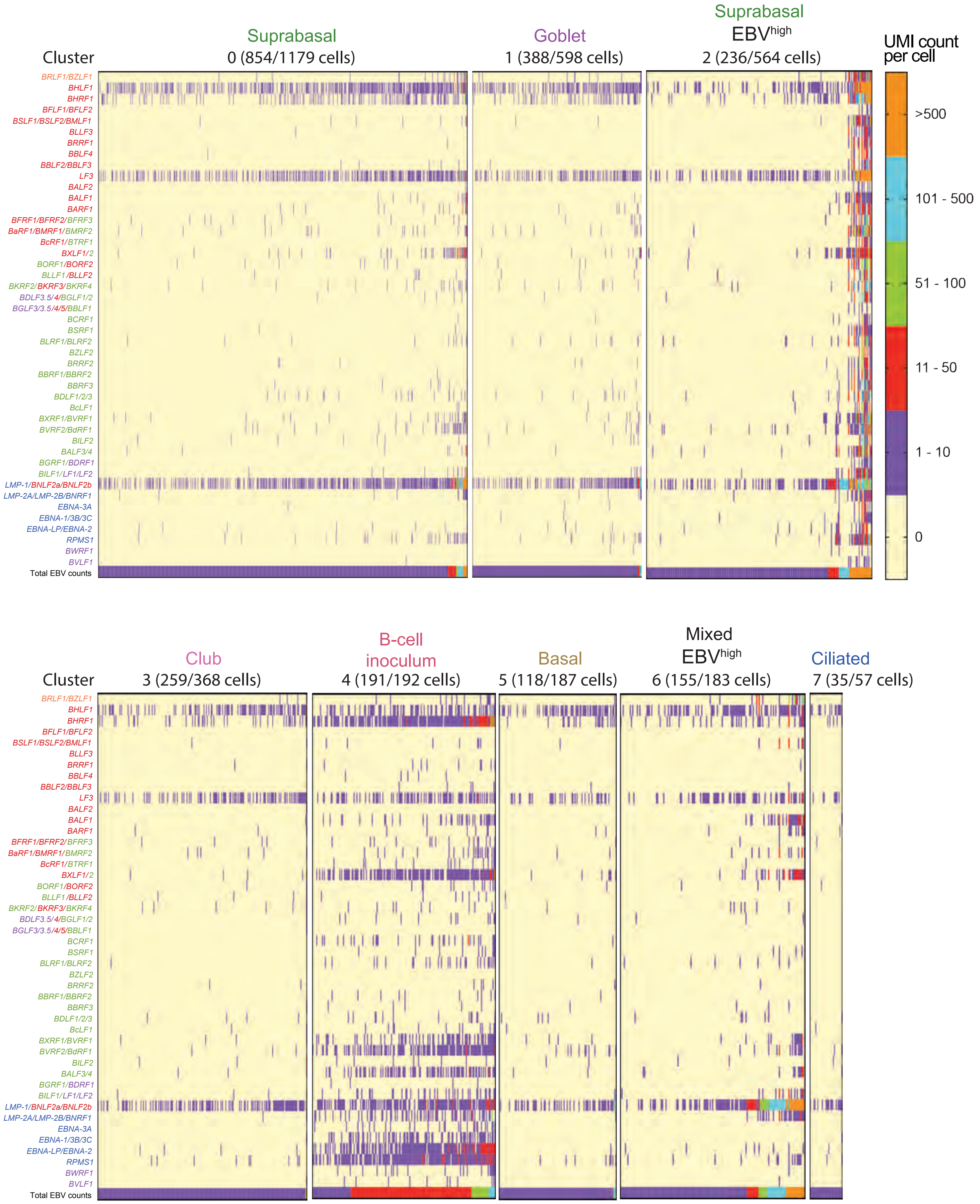


**Supplementary Figure 7. Distinguishing EBV gene expression profiles by cell type.** Heatmaps show EBV gene expression by UMI count per cell, displayed for each cluster, for the EBV-infected pseudo-ALI culture of donor no. 4. To distinguish EBV$^{high}$ from EBV$^{low}$ cells, total EBV counts (total EBV UMI counts per cell) are shown at the bottom of each heatmap. Only the EBV-infected cells are shown with the number of EBV-infected cells displayed for each cluster indicated in parenthesis. EBV genes are colour coded into lytic: immediate-early (orange)/early (red)/late (green); latent (blue); and unassigned (purple). The scRNA-seq reads were aligned to the EBV genome fused annotation.

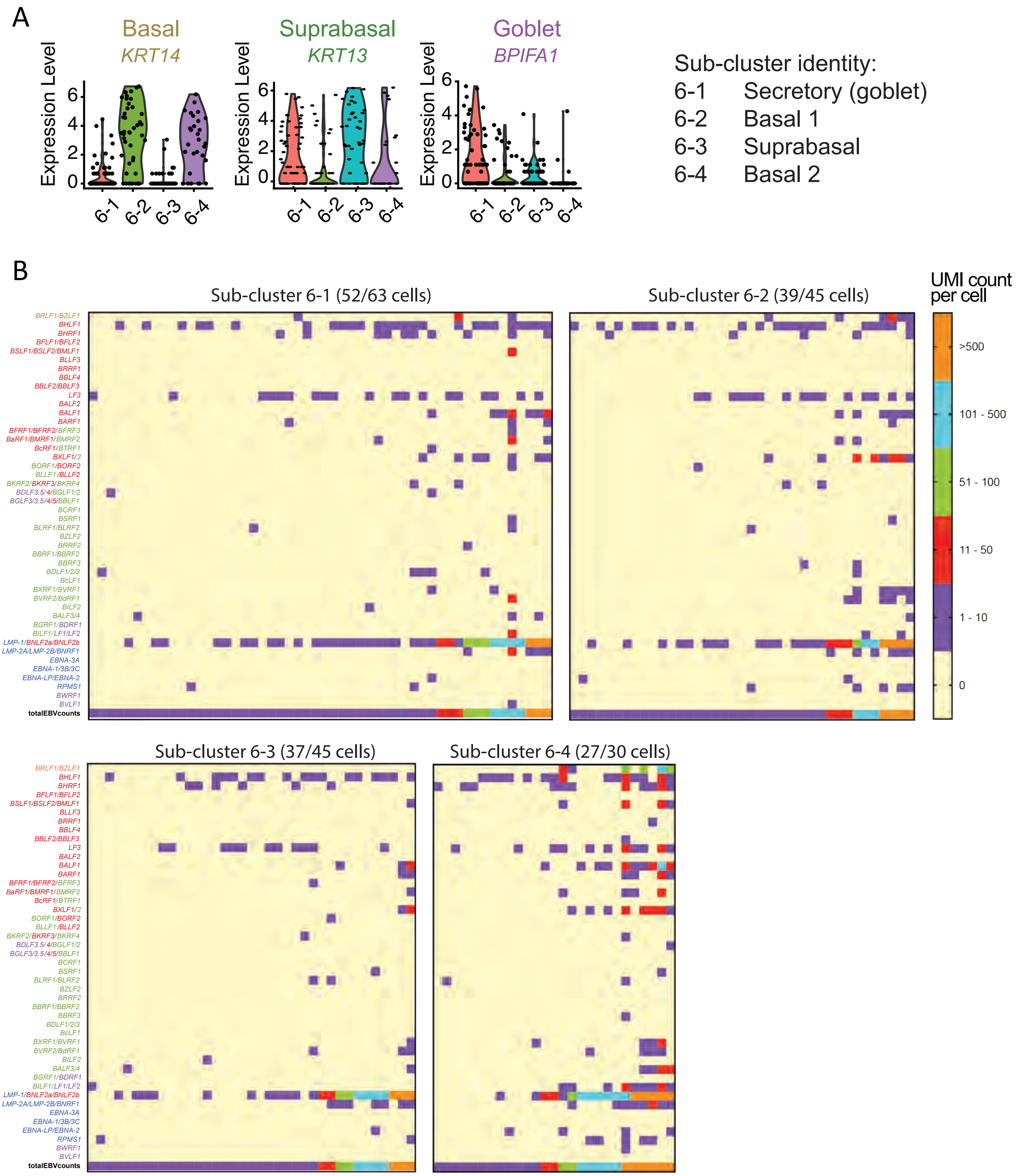


**Supplementary Figure 8. Separating the mixed cell cluster (cluster 6) by subclustering and EBV gene expression.** Shown are the results for the EBV-infected pseudo-ALI culture of donor no. 4. (A) Assignment of cell types by expression of marker genes for sub-clusters 6-1, 6-2, 6-3,and 6-4. (B) Heatmaps show EBV gene expression by UMI count per cell, displayed for each sub-cluster. $EBV^{high}$ cells can be distinguished from $EBV^{low}$ cells by the total EBV counts (total EBV UMI counts per cell) shown at the bottom of each heatmap. Only the EBV-infected cells are shown with the number of EBV-infected cells displayed for each sub-cluster indicated in parenthesis. EBV genes are colour coded into lytic: immediate-early (orange)/early (red)/late (green); latent (blue); and unassigned (purple). The scRNA-seq reads were aligned to the EBV genome fused annotation.

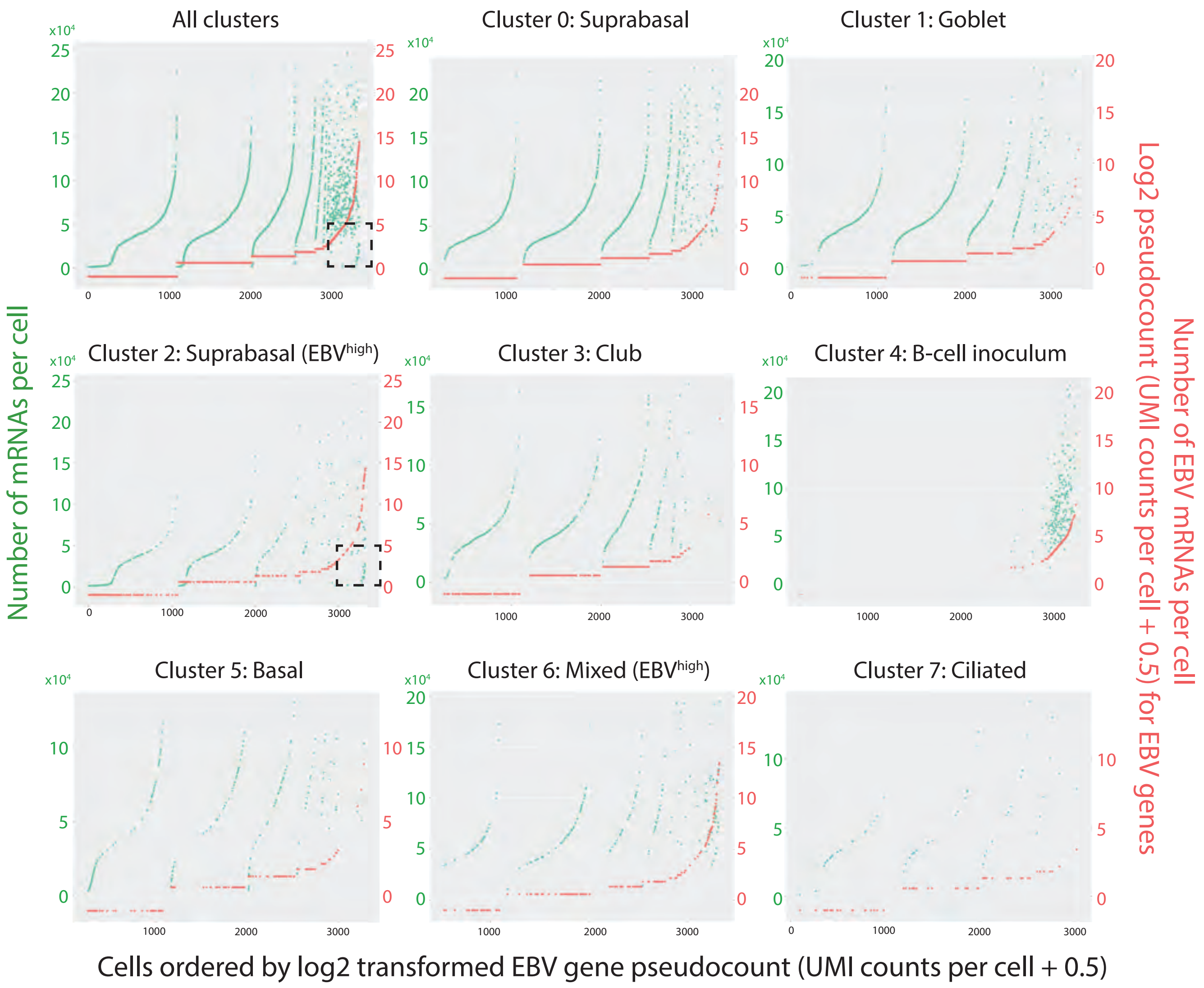


**Supplementary Figure 9. Comparison of EBV gene count to total number of mRNAs detected per cell.** Shown are the results for the EBV-infected pseudo-ALI culture from donor no. 4. Dot plots show EBV gene pseudocount (red) and the total number of mRNAs (green) per cell. Each dot represents one cell. Cells are ordered by the EBV gene pseudocount on the x-axes. Data are organized by cluster. Dashed black box, denotes cells with high numbers of EBV mRNAs but low total mRNA counts representative of host shut-off. Only cells with >1000 mRNAs per cell are shown as defined by the filtering criteria (see Supplementary Methods).

Supplementary Tables.

A. Summary of EBV qPCR and infectious titre results for pseudo-ALI cultures from donor no.4.

| Condition (days p.i.) | EBV DNA qPCR (n=3) | | | | | | Green Raji Unit [GRU] Titre (n=2) | |
|---|---|---|---|---|---|---|---|---|
| | Extracellular EBV genomes per ALI | | | Cell-associated EBV genomes per ALI | | | GRU per ALI | S.D. |
| | $C_T$ | Copy Number | S.D. | $C_T$ | Copy Number | S.D. | | |
| Day 2, rAkata-infected (n=2) | 28.84±0.08 | 3.13E+04 | 1.87E+03 | 28.56±0.12 | 4.06E+04 | 3.25E+03 | 624 | 238 |
| Day 5, rAkata-infected (n=6) | 23.90±0.07 | 1.16E+06 | 4.80E+04 | 30.23±0.31 | 1.55E+04 | 2.62E+03 | 1.07E+05 | 6.29E+04 |
| Day 2, Input Control (n=2) | 29.38±0.07 | 2.16E+04 | 1.27E+03 | 30.08±0.12 | 1.21E+04 | 1.28E+03 | 0 | 0 |
| Day 5, Input Control (n=3) | 29.46±0.15 | 2.00E+04 | 2.44E+03 | 31.40±0.34 | 4.67E+03 | 1.58E+03 | 8 | 8 |
| No Template or Media Control | 33.58±0.30 | 0 | 0 | 32.06±0.13 | 0 | 0 | 0 | 0 |

B. Summary of antibodies and staining reagents for image analysis.

| Stain | Target | Antibody (Ab)/Probe/Reagent | Antibody/Probe/Reagent Name | Concentration | Vendor | Part Number |
|---|---|---|---|---|---|---|
| 1 | Zebra | Primary Ab | Mouse anti-Zebra (clone BZ1) | 2 µg/mL | Santa Cruz Biotechnology | sc-53904 |
| | | Seconday Ab | Donkey anti-mouse IgG, Cy3 | 1 µg/mL | Jackson Immunoresearch | 715-165-150 |
| 2 | LMP1 | Primary Ab | Mouse anti-LMP1 (clone CS1-4) | 10 µg/mL | Abcam | ab78113 |
| | | Seconday Ab | Donkey anti-mouse Cy3 | 1 µg/mL | Jackson Immunoresearch | 715-165-150 |
| 3 | gp350 | Primary Ab | Mouse anti-gp350 (clone 0221) | 1 µg/mL | Santa Cruz Biotechnology | sc-57724 |
| | | Seconday Ab | Donkey anti-mouse IgG, Biotin-SP | 1 µg/mL | Jackson Immunoresearch | NC9673676 |
| | | Tertiary/Development Reagent | Donkey anti-mouse IgG, Biotin-SP | 1 µg/mL | Jackson Immunoresearch | NC9673676 |
| 4 | Human IgG | Primary Ab | Goat anti-human IgG, A647 | 2 µg/mL | Invitrogen | A-21445 |
| 5 | EphA2 | Primary Ab | Rabbit anti-EphA2 (clone D4A2) | 0.45 µg/mL | Cell Signaling Technologies | 6997S |
| | | Seconday Ab | SignalStain Boost Reagent (HRP, rabbit) | Undiluted | Cell Signaling Technologies | 8114S |
| | | Development Reagent | DAB | As recommended | Zytovision | T-1063-40 |
| 6 | EBER | EBER probe | ZytoFAST EBER Biotin Probe | Undiluted | Zytovision | T-1014-400 |
| | | Conjugate | Streptavidin-Rhodamine Red-X | 2 µg/mL | Jackson Immunoresearch | 016-290-084 |
