## Supplementary Materials for "A three-dimensional Air-Liquid Interface Culture Model for the Study of Epstein-Barr virus Infection in the Nasopharynx"

Supplementary Methods.

Supplementary Figures 1-9.

Supplementary Tables.

**SUPPLEMENTARY METHODS**

**Analysis of Virus Production**. Pseudo-ALI cultures were harvested 2 days p.i. (immediately after removal of B-cells and 3 HBSS washes) or 5 days p.i. by scraping in 100 μL total DPBS. Cells were pelleted by centrifugation at 500 xg for 5 min. Supernatant was removed and collected as extracellular virus. Cells were lysed by repeated freeze thawing, pelleted by centrifugation at >20000 xg for 10 min, and supernatant collected as cell-associated virus. To degrade unencapsidated cell-free DNA, supernatant was treated with 2 units of Turbo DNase (Thermo Scientific) per 50 μL supernatant for 30 mins at 37^o^C followed by inactivation for 10 mins at 70^o^C, and 100 μg/mL proteinase K (Fisher Scientific) for 30 mins at 50^o^C to release encapsidated DNA followed by inactivation at 75^o^C for 20 mins. EBV genomes were quantified by amplifying a region of *BALF5* using 0.3 μM each of primers with the sequences: 5′ GAGCGATCTTGGCAATCTCT 3′ and 5′ TGGTCATGGATCTGCTAAACC 3′. Quantitative PCR reactions were assembled using the Maxima SYBR Green qPCR master mix kit (Thermo Scientific) as per the manufacturer’s instructions, with 2 μL of supernatant containing template DNA, ROX as a passive reference dye, and absolute quantitation against a standard curve. Reactions were performed on the QuantStudio 3 instrument (Applied Biosystems) and analyzed with QuantStudio Design and Analysis Software (v1.4.1). Mean values and standard deviation were calculated from technical triplicates. A standard curve was generated using 25 to 10^6^ copies of a plasmid containing a fragment of *BALF5*. Harvested supernatant was titrated by the Green Raji Unit (GRU) method as previously described (*1*).

**Immunofluorescence, Immunohistochemistry and Image Analysis.** Immunostaining of EBV proteins was performed on whole mount transwell membranes as previously described (*1*). EBV-encoded RNAs (EBERs) were detected by fluorescent in-situ hybridization as per the manufacturer’s protocol (Zytovision). Detection and post hybridization steps were performed according to the immunofluorescence protein staining protocol. Primary antibodies, concentrations, and detection reagents are listed in Supplementary Tables B. Confocal images were captured on an Olympus IX81 inverted microscope, CREST X-Light V2 spinning disc confocal, with a 20x (numerical aperture [NA] = 0.45), 60x oil (NA = 1.45), or 100x oil (NA = 1.4) objective, and a Hamamatsu Flash 4.0 LT camera. Maximum intensity projections of z-stack images of ALI cultures (z spacing = 170 nm) were generated using Olympus CellSens Dimension software. Histograms comparing intensity of staining for EBV markers were generated by combining the 580 nm channel of the mock image and infected image in CellSens Dimension. LMP1 particle analysis was performed using Fiji software (https://imagej.net/Fiji). Particle intensity, circularity and size thresholds were applied to the red fluorescent channel. The Fiji plugin, analyze particles, was used to determine intensity of particles with parameters of 0-1 circularity and 0-0.001 pixel^2^ size. For histology, whole transwell membranes were fixed overnight in 4% paraformaldehyde. Paraffin embedding, Hematoxylin & eosin (H&E), Alcian blue/periodic acid Schiff, pan-cytokeratin (Dako, clone AE1/AE3), and Ki67 (Dako, clone MIB-1) staining were performed by the University of Pittsburgh Research Histology Services. Immunohistochemistry for Ephrin receptor A2 (Cell Signaling Technology, clone D4A2) was performed as per the manufacturer’s protocol with the following modification: ZytoVision 3,3′-Diaminobenzidine (DAB) reagent was used for development and nuclei were counterstained with ZytoVision Nuclear Blue. Brightfield images were captured on an Olympus PROVIS AX70 microscope with a QImaging QIClick camera and QCapture Pro 7 software.

**scRNA-seq analysis.** The 10X Genomics Cell Ranger pipeline for scRNA-seq counts unique molecular identifiers (UMIs) that are confidently mapped to one exonic locus of an annotated gene in a strand-specific manner. Unannotated, non-exonic reads and reads that map to loci of overlapping genes on the same strand are not counted. Sequencing reads were aligned to a merged human and EBV (Akata strain) genome (hg38+EBV) compiled from NCBI GenBank GRCh38.90 and KC207813.1 using the 10X Genomics Cell Ranger 3.0.2 workflow. Exon coordinates were added to the EBV genome using available information from annotated CDS, and when unavailable (*BSLF1, BBLF2/BBLF3, BALF3*) gene coordinates were used. Sequence coverage containing downstream complementary polyadenylation signal for *BHLF1* and *LMP-1/BNLF2a/BNLF2b* was observed and the annotations for the genes were extended accordingly. Seventy five EBV gene annotations were added to the original 13. The gff3 files were converted to gtf using gffread from cufflinks/2.2.1. An index of the merged genomes was created with cellranger/3.0.2 mkref. The Fastq files were analyzed with cellranger/3.0.2 count applying the SC3Pv3 chemistry. More than 470M reads were captured (sequencing saturation similar across clusters) with 123,249 mean reads per cell and 71.8% reads mapped confidently to the (EBV+hg38) transcriptome. Seurat (v3.1.4), an R package developed for single cell analysis, was used for data analysis, normalization of gene expression and visualization of cell populations (*2*). Cells were clustered using shared nearest neighbor modularity by the K-Nearest Neighbor algorithm implemented in Seurat. Cells were filtered for the following inclusion criteria: total number of detected genes > 500, percentage of mitochondrial genes < 25%, percentage of hemoglobin genes < 0.025%, percentage of ribosomal genes < 40%, and total number of read counts > 1000.

Supplementary Tables. (A) Summary of EBV qPCR and infectious titre results for pseudo-ALI cultures from donor no. 4. (B) Summary of antibodies and staining reagents for image analysis.

**SUPPLEMENTARY REFERENCES**

1.	E. A. Caves *et al.*, Air-Liquid Interface Method To Study Epstein-Barr Virus Pathogenesis in Nasopharyngeal Epithelial Cells. *mSphere* **3**, (2018).

2.	R. Satija, J. A. Farrell, D. Gennert, A. F. Schier, A. Regev, Spatial reconstruction of single-cell gene expression data. *Nature biotechnology* **33**, 495-502 (2015).
